# Model-informed Deep Q-Networks to Guide Infliximab Dosing in Pediatric Crohn’s Disease

**DOI:** 10.1101/2025.07.14.664804

**Authors:** Kei Irie, Phillip Minar, Jack Reifenberg, Brendan M Boyle, Joshua D Noe, S Jeffrey Hyams, Tomoyuki Mizuno

**Affiliations:** Division of Translational and Clinical Pharmacology, Cincinnati Children’s Hospital Medical Center, Cincinnati, Ohio, USA; Division of Gastroenterology, Hepatology and Nutrition, Cincinnati Children’s Hospital Medical Center, Cincinnati, Ohio, USA; Department of Pediatrics, University of Cincinnati College of Medicine, Cincinnati, Ohio, USA; University of Cincinnati School of Medicine, Cincinnati, Ohio, USA; Division of Gastroenterology, Hepatology and Nutrition, Nationwide Children’s Hospital, Columbus, Ohio, USA; Division of Gastroenterology, Hepatology and Nutrition, Children’s Hospital of Wisconsin, Milwaukee, Wisconsin, USA; Division of Gastroenterology, Hepatology and Nutrition, Connecticut Children’s Medical Center, Hartford, Connecticut, USA

**Keywords:** Model-informed precision dosing, Reinforcement learning, Deep learning, Children, Inflammatory bowel disease

## Abstract

Model-informed precision dosing (MIPD) utilizes pharmacokinetic/pharmacodynamic (PK/PD) models to optimize drug therapy. However, conventional MIPD often requires manual simulation and regimen selection, which are time-consuming and demand specialized expertise. Reinforcement learning (RL), in which an agent learns optimal decisions through iterative interactions with an environment, offers a scalable and automated alternative. In this study, we developed a model-informed Deep Q-Network (DQN) to personalize infliximab dosing for patients with Crohn’s disease. The DQN was trained in a simulation environment incorporating a population PK model, inter-individual variability, and assay error. Virtual patients with randomly sampled covariates were used to explore dosing strategies at infusions 1, 3, and 4. Doses ranged from 1 to 10 mg/kg at infusion 1 and from 1 to 20 mg/kg thereafter, with intervals of 4 to 12 weeks. The reward function prioritized achieving trough concentrations of 18–26 µg/mL before infusion 3 and 5–10 µg/mL before infusions 4 and 5, while penalizing overtreatment and additional infusions. The DQN policy converged after 80,000 episodes, yielding target attainment probabilities (PTAs) of 92.9% and 98.4% at infusions 4 and 5, respectively, in 1,000 virtual patients. High doses (11–20 mg/kg) were selected in only 0.2% of cases. At infusion 4, 66.8% of patients received an 8-week interval, and 57.3% at infusion 5. Retrospective real-world validation showed that patients whose actual doses matched DQN recommendations had trough levels significantly closer to target ranges. These findings support the feasibility of using DQN-based agents to enhance and automate infliximab individualized dosing in pediatric populations.

## INTRODUCTION

Infliximab, a monoclonal antibody targeting tumor necrosis factor-alpha (TNF-α), is a cornerstone therapy for children with moderate to severe Crohn’s disease (1). Maintaining optimal infliximab exposures is essential for achieving sustained therapeutic efficacy and long-term disease remission (2). A trough concentration range of >5 μg/mL during maintenance therapy improves disease remission rates and enhances durability of the anti-TNF biologics (2). Additionally, infliximab concentration at infusion 3 (week 6) ≥18 μg/ml is associated with early clinical and biological responses (3). However, standard weight-based dosing (5 mg/kg) often fails to achieve these therapeutic trough concentrations in pediatric patients. Substantial inter-individual pharmacokinetic (PK) variability poses a major challenge to achieving consistent infliximab exposure. Key influencing factors to infliximab clearance include body weight (WT), albumin levels (ALB), erythrocyte sedimentation rate (ESR), neutrophil CD64 (nCD64), and anti-infliximab antibodies (ATI) (4).

Therapeutic drug monitoring (TDM) and model-informed precision dosing (MIPD) are key strategies for personalizing infliximab therapy by integrating patient-specific factors with population PK models (5). While these approaches offer personalized dosing strategies, the process typically involves several steps, including performing simulations with population PK models and exploring the dosing regimens (i.e., dose amount and interval) to achieve pre-defined target concentrations. This individualized, case-by-case process is often time-consuming, causing potential variations among performers, and challenging to implement consistently across clinical settings. With the increasing adoption of MIPD in clinical practice, there is a growing need for automated and standardized approaches to support timely, reproducible, and scalable dosing decisions. Recent technological advances, including digital dashboards integrated into clinical workflows, are beginning to address this need by enabling real-time, patient-specific dosing recommendations (6).

Reinforcement learning (RL) has recently emerged as a promising approach to enhance precision dosing strategies (7–10). Although RL is ideally suited for adaptive dosing, it is theoretically challenging to train algorithms solely on data from real-world patients, as it typically lacks sufficient cases with suboptimal or unsafe clinical conditions (10). The MIPD framework can address this challenge by utilizing mechanistic population PK/PD models and virtual patient cohorts, simulating diverse dosing scenarios without patient risk. Combining RL with model-informed simulations thus enables RL algorithms to be trained in safe, data-rich environments that account for individual variability and facilitate reward-based decision-making. Q-learning and its deep-learning extension, deep Q-networks (DQN), have been successfully applied to optimize dosing regimens for several drugs, including anticancer agents (11) and warfarin (12, 13). These studies highlight the potential of model-informed RL, combined with population PK/PD models, to refine dose selection. While RL is being explored for dosing in various therapeutic areas, biologics used in pediatrics, such as infliximab, have not yet been studied using DQN approaches.

In this study, we developed an online DQN framework to optimize infliximab dosing in pediatric patients with Crohn’s disease. The DQN model was designed to interact with a simulation environment, integrating a population PK model and virtual pediatric patients. Incorporating proactive TDM scenarios and key covariate information influencing infliximab clearance, the model identified individualized dosing strategies to achieve target trough concentrations. This approach, integrating MIPD and RL, offers automated and standardized dose selection of infliximab in pediatric clinical care.

## METHODS

### Q-Learning Framework for Optimal Infliximab Dosing Strategy

Q-learning was employed to identify optimal infliximab dosing regimens in virtual pediatric patients, as illustrated in **Figure 1**. Virtual patients were generated by randomly sampling key covariates (WT, ALB, ESR, nCD64, and ATI) from log-normal distributions derived from real-world data (4). The established pediatric population PK model (4, 14) was implemented for PK simulations as described in the **Supplemental methods**.

**Figure 1.**
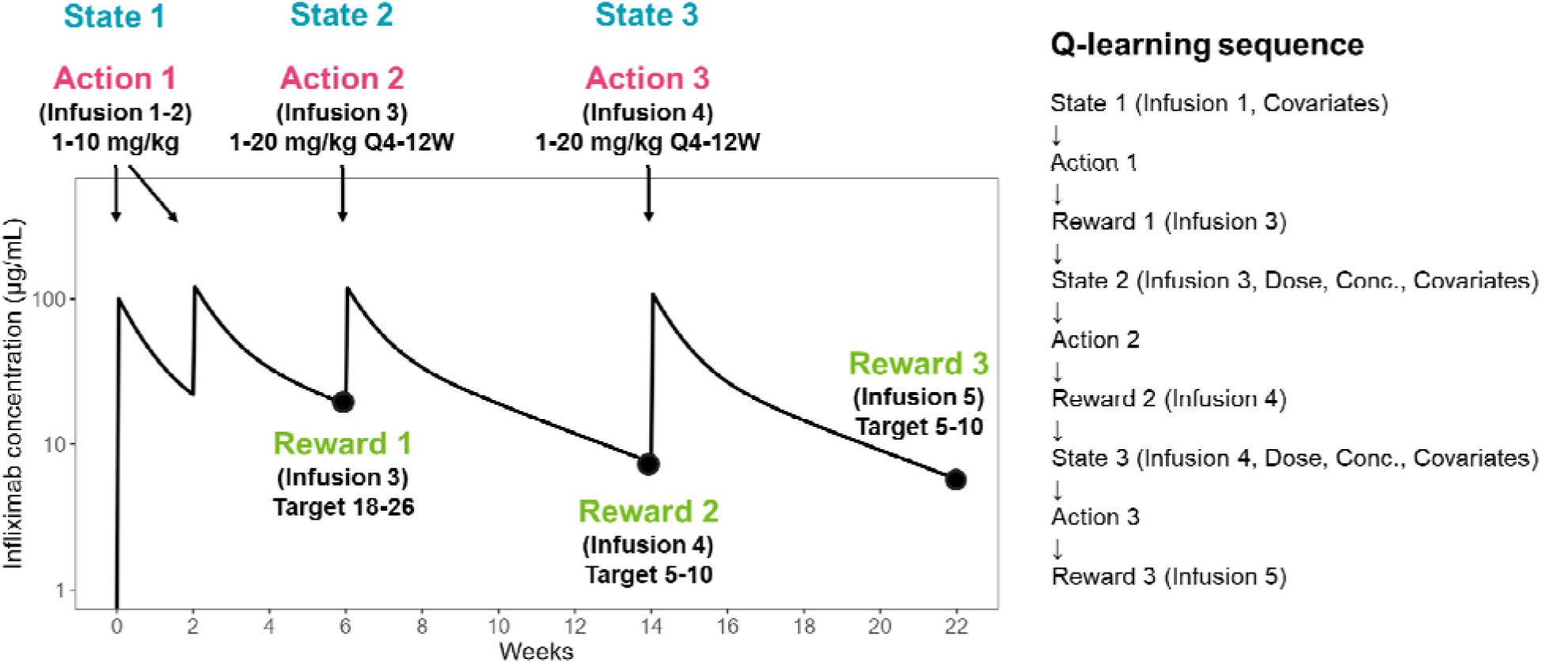
Q-learning sequence for selecting the optimal infliximab dosing regimens.

At infusion 1 (State 1), an initial dose (Action 1) ranging from 1 to 10 mg/kg was selected by the algorithm based on patient-specific covariates. The same dose was applied at infusion 2 (week 2), and the plasma concentration-time profile for the selected dose was subsequently simulated. A reward was then calculated based on the simulated trough concentration at week 6 (infusion 3) by assuming that the first proactive TDM was performed before infusion 3 (State 2), based on evidence supporting early TDM in IBD (15, 16). The second dosing decision (Action 2) was made using an updated state that included the initial dose, the week 6 trough concentration, and the covariate data. Doses at this stage ranged from 1 to 20 mg/kg and were administered at intervals of 4 to 12 weeks. The PK profile was again simulated, and the reward was determined based on the resulting trough concentration at infusion 4 (week 10–18, depending on the interval selected by the agent). At infusion 4 (State 3), a third dosing decision (Action 3) was determined using a further updated state that incorporated the doses from infusions 1–3, the trough concentration at infusion 4, and patient covariates. The final reward was based on the simulated trough concentration at the infusion 5 (week 14–30, based on the selected interval).

During training, the Q-learning algorithm explored a wide range of dosing strategies at each decision point, simulating outcomes to maximize expected rewards. The reward (*R*) wa defined as the negative absolute difference between the resulting trough concentration (*C*) and the target median concentration, calculated as:

Where Target_median_ was set to 21 µg/mL for infusion 3 (target range: 18–24 µg/mL) and 7.5 µg/mL for infusions 4 and 5 (target range: 5–10 µg/mL) (2, 3).

To account for the priority of dosing strategies, an additional decision-based reward function was evaluated as follows:

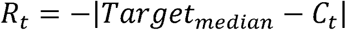

- If *C* is within the target range, then *R + 3*
- If *C* is within the target range and the dose is within 1–10 mg/kg, then *R + 6*
- If *C* is within the target range, the dose is 1–10 mg/kg, and the dosing interval is every 8 weeks, then *R + 9*

These reward structures were designed to discourage higher doses (11–20 mg/kg) when lower doses (1–10 mg/kg) were sufficient to achieve target concentrations. Additionally, a standard 8-week dosing interval was preferred to minimize the number of clinic visits when adequate drug exposure could be maintained. This design guided the model toward dosing strategies that were both clinically effective and operationally practical. Detailed definitions of the state space, action space, and Q-value formulation are provided in the **Supplemental Methods**.

### Development of Model-informed Deep Q-Networks

Given the continuous nature of patient-specific covariates and drug concentrations, which results in a high-dimensional and potentially infinite state space, an online double DQN with a dueling network architecture and a prioritized experience replay buffer (17–20) was employed to approximate the Q-function. **Figure 2** illustrates the framework for online model-informed DQN training. The agent was trained within a simulation environment that combined the population PK model with virtual pediatric patients. In each episode, the agent observed the current state and selected a dose (action). The PK response (i.e., concentration-time profile) was then simulated, the state was updated based on the resulting concentration, and a reward was assigned according to the simulated trough concentration. This process was repeated for infusions 1, 3, and 4. Upon completing all decision points, the episode concluded, and a new virtual patient was generated for the next training iteration. Each experience tuple (state, action, reward, and next state) was stored in a replay buffer. Once the number of stored experiences exceeded the batch size, mini-batches were randomly sampled for training. Target Q-values were computed based on the observed rewards and future Q-values estimated by a separate target network, which was synchronized with the main network to stabilize DQN training. The neural network was trained by minimizing the Huber loss between the predicted and target Q-values (17). This learning process was repeated for 80,000 episodes. The detailed implementation of DQN is provided in the **Supplemental methods**.

**Figure 2.**
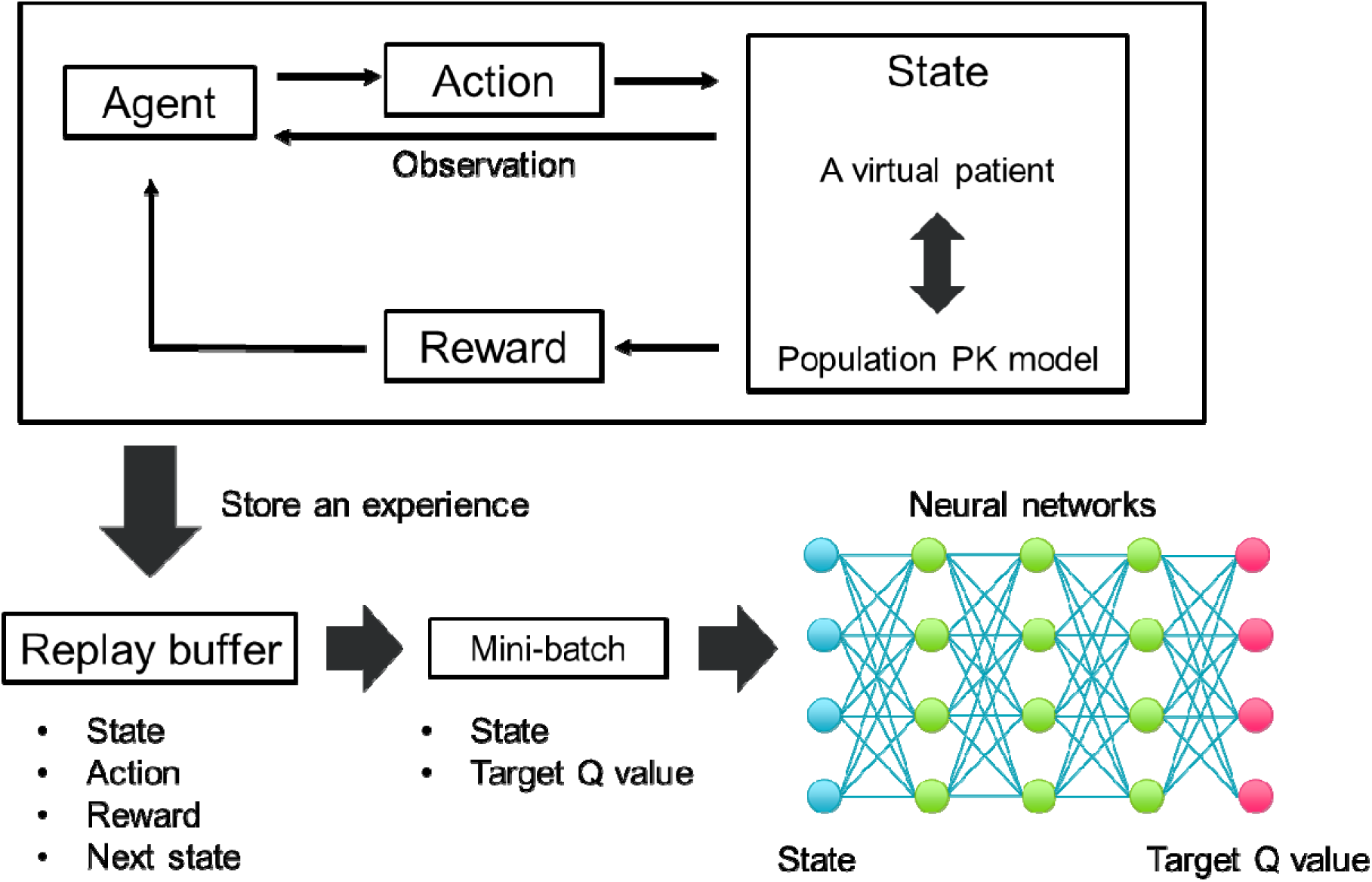
Online Model-Informed Deep Q-Network (MI-DQN) Framework. In each episode, the agent observed the current state, interacted with a PK simulation using virtual pediatric patients, and selected a dose strategy. The resulting experience—state, action, reward, and next state—was stored in a replay buffer. Once sufficient experiences were accumulated, mini-batches were sampled to train the Q-network by minimizing the loss between predicted and target Q-values.

### Evaluation of Deep Q-Networks framework

To benchmark the DQN framework, models were sequentially trained and evaluated under progressively complex simulation conditions:

1. A PK model without inter-individual variability (IIV) and assay error (i.e., virtual patients with PK varied solely by covariates).
2. A PK model incorporating IIV (25.1%) in CL (4).
3. A PK model incorporating both IIV in CL and 10% of assay error (21).
4. A PK model with IIV in CL, 10% of assay error, and a decision-based reward function.

Models 1–3 were designed to evaluate the stability and robustness of the DQN under progressively increasing levels of biological variability and measurement noise. Model 4 introduced a decision-based reward function designed to promote clinically preferred dosing behaviors. By integrating these practical considerations into the reward structure, the model learned to avoid unnecessary dose escalation when lower doses were sufficient, while also favoring an 8-week interval to align with the current standard dosing scheme.

Model performance was assessed during training by tracking the probabilities of target attainment (PTA) at infusions 3, 4, and 5. These evaluations were conducted every 100 training episodes using an independent cohort of 1,000 virtual pediatric patients. Final PTA outcomes under DQN-guided dosing were compared to those achieved using a conventional fixed dosing regimen of 5 mg/kg. To ensure robustness and generalizability, comparisons were averaged across 10 distinct cohorts of 1,000 virtual patients, each generated with a different random seed.

### Evaluation of Deep Q-Networks with real-world data

The predictive performance of the DQN was retrospectively evaluated using real-world data from two clinical studies involving pediatric and young adult patients aged 2 to 22 years receiving infliximab for Crohn’s disease. The first study, REFINE, was a multicenter, prospective observational study conducted at Cincinnati Children’s Hospital Medical Center (CCHMC), Connecticut Children’s Medical Center, the Medical College of Wisconsin, and Nationwide Children’s Hospital between 2014 and 2019. The second study, APPDASH, enrolled patients at CCHMC between 2019 and 2021. Both studies were approved by their respective Institutional Review Boards.

Patients were included if they had complete covariate data and received dosing at week 0, week 2 ± 3 days, week 6 ± 3 days, week 10-18 ± 3 days, and week 14-30 ± 3 days. For evaluation of the initial dosing decision, patients with available infliximab trough concentrations at infusion 3 were included. For evaluation of subsequent dosing decisions, patients with available concentrations at both infusion 3 and 4, or infusion 4 and 5, were included.

The negative absolute difference between the observed and target concentrations (nAD_Target_) was compared to the negative absolute difference between the actual administered dose or dose per intervals and the DQN-predicted dose or dose per intervals (nAD_DQNdose_) defined as follows:

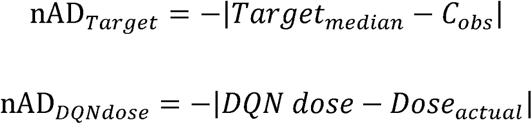

Where Target_median_ was set to 21 µg/mL for infusion 3 and 7.5 µg/mL for infusions 4 and 5. The observed trough concentration (µg/mL) was denoted as C_obs_. The DQN dose refers to the dose (mg/kg) or dose per interval (mg/kg/week) predicted by the DQN based on the patient’s clinical state, while Dose_actual_ represents the actual administered dose (mg/kg) or dose per interval (mg/kg/week).

The negative absolute difference from the target concentration, denoted as nAD_Target_, reflects how closely the observed level aligned with the target. Similarly, the negative absolute difference from the DQN prediction, denoted as nAD_DQNdose_, reflects how closely the actual dose aligned with the DQN-recommended dose. In both cases, values closer to zero indicate better alignment with the target or DQN recommendation. Correlation between nAD_Target_ and nAD_DQNdose_ was evaluated using Spearman’s rank correlation test, with p-values < 0.05 considered statistically significant.

## RESULTS

### Development of Deep Q-Networks (DQN)

The trajectory of the PTA during the DQN training in 1,000 virtual patients under four simulation conditions is shown in **Figure 3**. The PTA for pre-dose concentrations at infusions 3, 4, and 5 gradually stabilized over 80,000 training episodes, indicating that the DQN models progressively learned an effective dosing strategy and successfully converged. The PTA was lower at infusion 3 compared to infusions 4 and 5, primarily due to the restricted initial dose range (1–10 mg/kg) and IIV in CL, which could not be fully predicted from covariates available at infusion 1. However, the dosing strategy was refined based on observed drug concentrations, resulting in 100% PTA at infusions 4 and 5 under conditions without IIV (**Figure 3A**) and with IIV (**Figure 3B**). These results highlight the potential of the DQN framework to theoretically achieve perfect target attainment when dosing decisions are fully informed by reliable, noise-free data (covariates and observed concentrations). When assay errors were incorporated into the observed concentrations during simulations (**Figures 3C and 3D**), the PTA decreased slightly. However, the model maintained nearly 100% PTA at infusion 5, demonstrating the robust performance even when considering random assay variability. The resulting mean trough concentrations ± standard deviations were 18.4 ± 7.9 µg/mL, 7.2 ± 1.4 µg/mL, and 7.3 ± 1.0 µg/mL at infusions 3, 4, and 5, respectively, in the model incorporating IIV, assay error, and a decision-based reward function.

**Figure 3.**
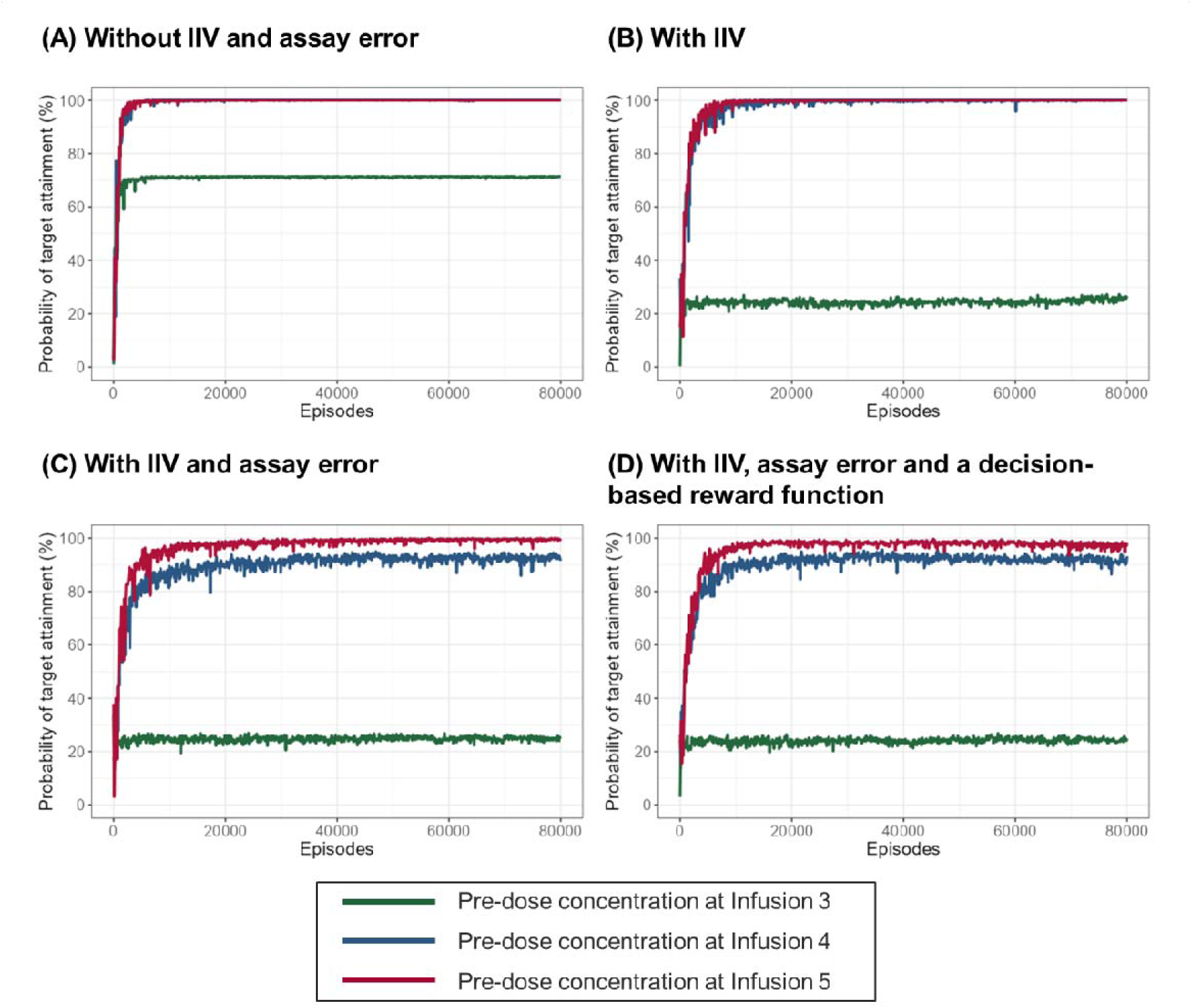
The trajectories of the PTA at infusion 3, 4, and 5 in 1,000 virtual patients under four simulation conditions. (A) A PK model without IIV and assay error (PK behavior of virtual patients varied solely by covariates). (B) A PK model incorporating IIV in CL. (C) A PK model incorporating both IIV in CL and assay error. (D) A PK model with IIV in CL, assay error, and a decision-based reward function.

**Table 1** summarizes the dosing strategies with and without decision-based reward functions, which were designed to discourage unnecessary dose escalation and reduce the burden of additional clinic visits. With this reward structure, high doses (11–20 mg/kg) were selected in only 0.2% of virtual patients at infusions 4 and 5, while the standard 8-week interval remained feasible for 66.8% and 57.3% of patients at infusions 4 and 5, respectively. These findings suggest that the DQN effectively integrated clinically relevant decision-making through reward function optimization. Additionally, PTA outcomes were consistent across 10 random seeds used for PK simulations and patient generation, consistently outperforming standard fixed dosing of 5 mg/kg (**Table 2**). The model incorporating IIV, assay error, and a decision-based reward function yielded PTAs of 25.6% ± 1.0 at infusion 3, 92.9% ± 0.7 at infusion 4, and 98.4% ± 0.5 at infusion 5.

**Table 1.** Selection of dosing strategies by DQN with and without decision-based reward functions in 1,000 virtual patients.

| Dose | Without Decision-Based Reward |  |  | With Decision-Based Reward |  |  |
| --- | --- | --- | --- | --- | --- | --- |
|  | 1-10<br>mg/kg | 11-20<br>mg/kg | 1-10<br>mg/kg<br>and<br>8 weeks<br>interval | 1-10<br>mg/kg | 11-20<br>mg/kg | 1-10<br>mg/kg<br>and<br>8 weeks<br>interval |
| Infusion 1 | 100% | - | - | 100% | - | - |
| Infusion 3 | 58.7% | 41.3% | 3.4% | 99.8% | 0.2% | 66.8% |
| Infusion 4 | 67.8% | 32.2% | 0.9% | 99.8% | 0.2% | 57.3% |

**Table 2.** Summary of PTA outcomes with DQN-guided dosing and standard body weight-based dosing (5 mg/kg) in 1,000 virtual patients.

| Dose | Without IIV and assay error |  | With IIV |  | With IIV and assay error |  | With IIV, assay error and a decision-based reward function |  |
| --- | --- | --- | --- | --- | --- | --- | --- | --- |
|  | 5 mg/kg | DQN | 5 mg/kg | DQN | 5 mg/kg | DQN | 5 mg/kg | DQN |
| Infusion 3 | 11.3%±0.9 | 72.9%±1.1 | 13.9%±1.3 | 25.5%±0.7 | 13.9%±1.3 | 25.5%±0.7 | 13.9%±1.3 | 25.6%±1.0 |
| Infusion 4 | 31.9%±1.6 | 100%±0 | 27.8%±1.5 | 100%±0 | 27.8%±1.5 | 94.7%±0.3 | 27.8%±1.5 | 92.9%±0.7 |
| Infusion 5 | 14.5%±1.2 | 100%±0 | 22.4%±1.8 | 100%±0 | 22.4%±1.8 | 99.4%±0.2 | 22.4%±1.8 | 98.4%±0.5 |

### Representative Virtual Patient Cases

Figure 4 presents two representative virtual patient cases to illustrate how the DQN model functions in individualized dosing. The model trained with IIV in CL, assay error, and a decision-based reward function was used. In Case 1, the patient was characterized by relatively low body WT and low ALB, along with significantly elevated ESR and nCD64, which are typically associated with increased CL and a need for higher dosing. The DQN model selected the maximum initial dose (10 mg/kg); however, the trough concentration fell below the target range at infusion 3. In response, the model intensified the dosing interval to every 5 weeks for infusion 3 and every 4 weeks for infusion 4, while still maintaining a dose range of 1–10 mg/kg. As a result, trough concentrations at infusions 4 and 5 were successfully maintained within the target range, demonstrating the model’s ability to avoid unnecessarily high dosing while maintaining efficacy.

**Figure 4.**
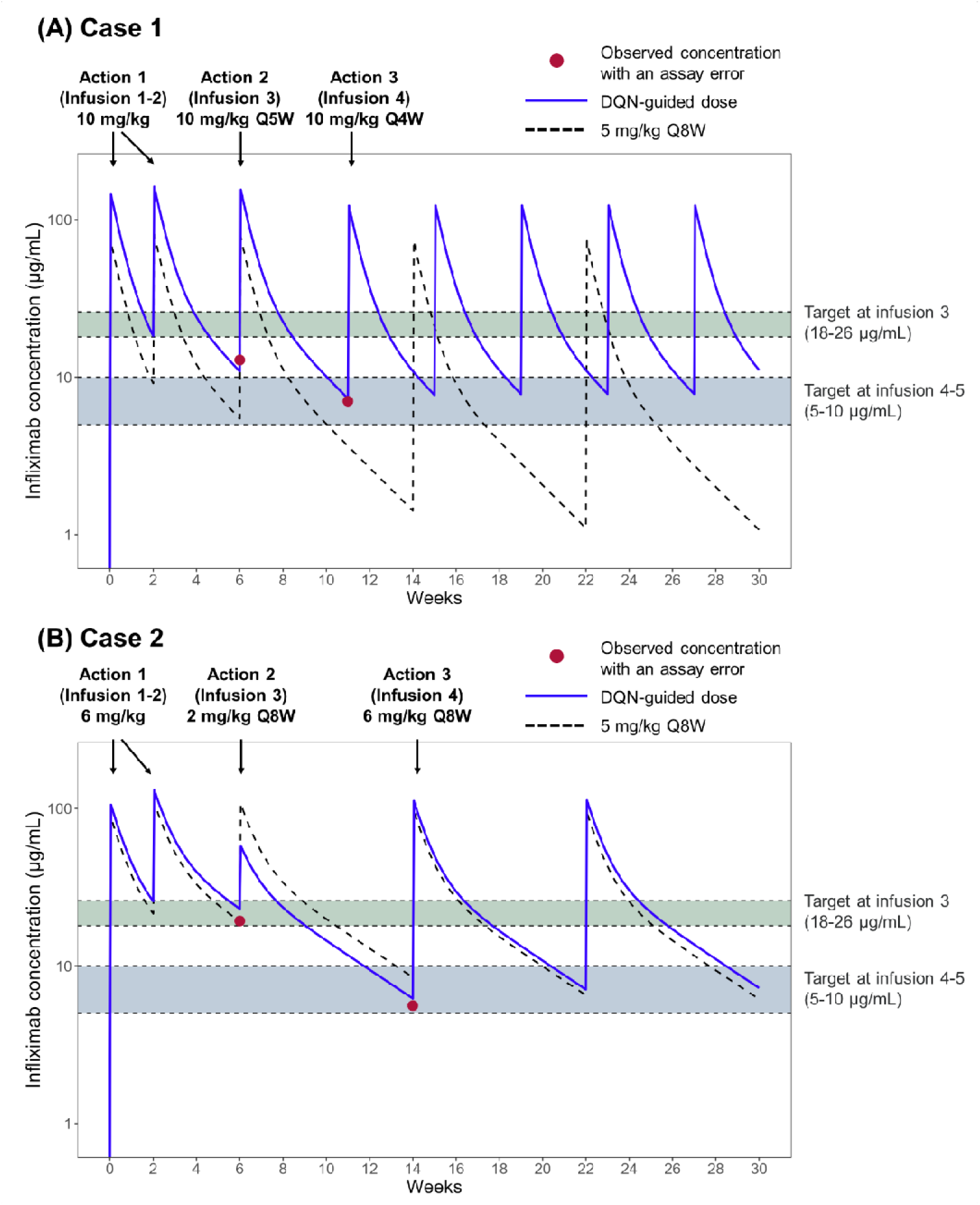
Representative Virtual Patient Cases in DQN-guided dose and standard dose. (A) Case 1 covariates: WT= 31.3 kg, ALB = 2.4 g/dL, ESR = 16.2 mm/h, nCD64 ratio = 20.5 ATI at infusion 3 = 36.0. (B) Case 2 covariates: WT= 46.6 kg, ALB = 3.7 g/dL, ESR = 3.9 mm/h, nCD64 ratio = 6.1, ATI at infusion 3 = 24.

**Figure 5.**
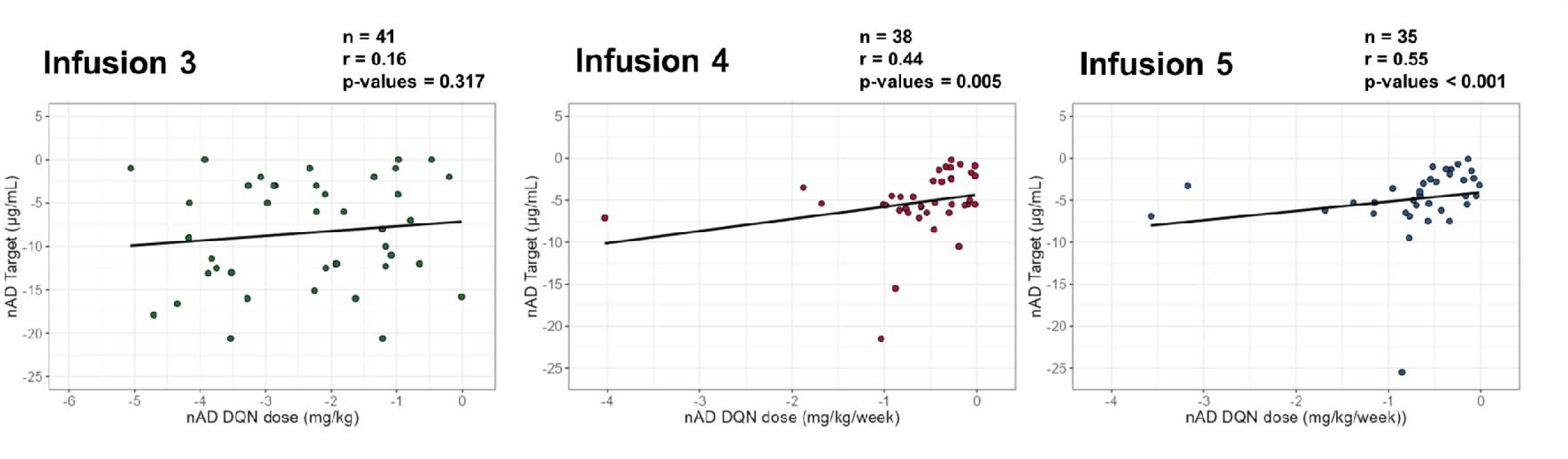
Correlation between nAD_Target_ (negative absolute difference from the target infliximab concentration) and nAD_DQNdose_ (negative absolute difference between the actual administered dose and the DQN-predicted dose).

In Case 2, an initial dose of 6 mg/kg was selected based on the patient’s relatively low inflammatory markers. The observed concentration at infusion 3 was within the target, however, it was affected by substantial assay error, yielding a measured value of 19.3 µg/mL, although the true concentration was below the target at 23.0 µg/mL. Despite this noise, the DQN model adjusted the dose to 2 mg/kg, which successfully maintained trough concentrations within the target range of 5–10 µg/mL, while preserving the standard 8-week dosing interval. This case highlights the model’s robustness in managing dosing decisions under conditions of assay variability.

### Evaluation of DQN-Guided Dosing with Real-World Data

The DQN-guided dosing strategy was retrospectively evaluated using real-world data from pediatric and young adult patients with Crohn’s disease treated with infliximab. The evaluation utilized the DQN model trained under simulations with IIV, assay errors, and decision-based reward functions, as this environment most closely reflects realistic clinical conditions. **Table S1** summarizes the characteristics of the patient cohorts included in the analysis. The negative absolute difference between observed and target infliximab concentrations (nAD_Target_) was compared to the negative absolute difference between the actual administered dose and the DQN-predicted dose (nAD_DQNdose_). No correlation was observed at infusion 3 as expected, likely due to the model’s limited predictive ability when using only baseline covariate data. However, a statistically significant correlation emerged at infusions 4 and 5, where dosing predictions incorporated observed concentrations. This suggests that patients receiving doses more closely aligned with DQN predictions were more likely to achieve target concentrations.

## DISCUSSION

In this study, we developed and evaluated a DQN framework to individualize infliximab dosing in pediatric Crohn’s disease, representing the first application of Q-learning to infliximab and its first validation using real-world data. Trained in a simulated environment incorporating a population PK model, IIV, assay error, and clinical dosing constraints, the DQN learned to optimize both induction and maintenance dosing to achieve target trough concentrations while minimizing overtreatment and preserving practical 8-week intervals. Retrospective analysis revealed that closer alignment with DQN-recommended doses was associated with improved proximity to target levels, supporting the potential of this approach as a scalable and automated method for precision dosing in pediatric care.

In this study, we implemented a Q-learning framework that mirrors proactive TDM processes conducted across the induction and maintenance phases of infliximab therapy. The agent sequentially observes the patient’s clinical state, including drug concentrations, covariates, and dosing history, and selects personalized dosing strategies to maximize pre-defined target attainment. This feedback-driven design aligns with clinical TDM, where dosing decisions are regularly adapted to the patient’s evolving status. Prior studies have demonstrated that Q-learning is well suited to such tasks. Anzabi Zadeh et al. (12) used deep Q-learning to optimize warfarin dosing based on observed INR values. De Carlo et al. (22) combined Q-learning with PK/PD modeling to personalize erdafitinib dosing, using serum phosphate as a biomarker to guide adaptive treatment. These examples highlight the potential of Q-learning to support biomarker-driven, state-based precision dosing.

The stepwise training process demonstrated the DQN model’s robustness under increasingly complex conditions. At the first infusion, the DQN based its decision solely on baseline covariates, reflecting population-level predictions similar to conventional MIPD (23–25). For later infusions, it updated decisions using observed concentrations—an approach conceptually similar to Bayesian prediction in conventional MIPD, which accounts for both IIV and assay error. By simulating assay noise during training, the DQN learned to make stable dosing decisions under realistic uncertainty. The reward function also promoted clinically preferred behaviors, such as avoiding unnecessary dose escalation and maintaining standard intervals, enabling consistent and interpretable recommendations. This framework supports the automation and standardization of complex dosing, reducing reliance on manual simulations and expert input, and paving the way for integration into digital health platforms. While model-informed RL has been applied in previous MIPD studies (11–13, 22, 26–28), real-world validation remains limited. This study demonstrated that greater concordance with DQN dosing in actual patients was significantly associated with better target concentration attainment, reinforcing its translational value. It is important to note, however, that this association also reflects the predictive accuracy of the underlying PK model, not just DQN itself. Incorporating additional covariates or biomarkers may help reduce uncertainty and enhance dosing precision. Continued refinement of both the PK model and input features may further unlock the potential of RL-based strategies across diverse patient populations.

The DQN framework supports broader efforts to scale and standardize MIPD (7, 10). By encoding expert-level decisions into a reproducible algorithm, it can reduce variability in clinical practice and support real-time, patient-specific recommendations, especially when embedded into electronic health record systems and clinical dashboards. Although further prospective clinical validation and algorithm refinement are necessary, the approach should be readily compatible with clinical decision support platforms such as RoadMAB (6), providing clinicians with automated, model-informed dosing suggestions during routine care. The DQN algorithm can rapidly evaluate a range of dosing strategies tailored to individual characteristics and propose optimal regimens aligned with therapeutic goals. This automated process reduces cognitive and operational burden, supports consistency in complex decision-making, and enables timely, personalized treatment adjustments within the clinical workflow. Incorporating patient- or institution-specific constraints could further strengthen its role in supporting nuanced and context-sensitive therapeutic decisions at the point of care.

Nonetheless, some limitations should be acknowledged. The performance of our DQN depends on the accuracy and generalizability of the PK model, which may not fully capture all relevant biological or disease-specific factors. This is particularly evident in the model’s limited ability to guide initial dose selection, which reflects its constrained capacity to predict PK in the absence of any drug concentration data. Improvements to the PK model, incorporating more predictive covariates, will be essential to enable more accurate and individualized decision-making in future iterations of the DQN framework. In addition, the limited PD biomarkers (e.g. fecal calprotectin or clinical remission status) incorporated into the PK model also limits the model’s ability to link exposure to outcomes.

In conclusion, this study represents a meaningful advance in precision dosing by being the first to apply and validate a Q-learning–based framework for infliximab. By enabling automated, standardized, and individualized dosing decisions, the DQN offers a promising solution to improve therapeutic outcomes and advance model-informed care in pediatric Crohn’s disease.

### What is the current knowledge on the topic?

Infliximab is a cornerstone therapy for pediatric Crohn’s disease; however, achieving therapeutic trough concentrations (5–10 µg/mL) is challenging due to high interindividual pharmacokinetic (PK) variability. Standard weight-based dosing is often suboptimal, and manual model-informed precision dosing (MIPD) is labor-intensive and can vary across clinicians.

### What question did this study address?

Can reinforcement learning (RL), specifically a Deep Q-Network (DQN), be used to automate and standardize individualized infliximab dosing decision-making processes in the treatment for pediatric Crohn’s disease? Can such a model trained in simulation be validated using real-world clinical data?

### What does this study add to our knowledge?

This is the first study to apply Q-learning to infliximab therapy and to validate an RL-based precision dosing framework using clinical data. The DQN identified effective dosing strategies in simulations, achieving >90% target attainment at later infusions while avoiding unnecessary high-dose use and maintaining standard 8-week intervals. Retrospective analysis showed that real-world patients whose dosing aligned with DQN recommendations were more likely to achieve target concentrations.

### How might this change clinical pharmacology or translational science?

This study demonstrates the potential of RL-guided MIPD as an automated, standardized, and scalable approach for biologic therapy. Embedding DQN frameworks into clinical dashboards may enable real-time, patient-specific dosing recommendations, improving therapeutic consistency, efficiency, and outcomes in pediatric patients.

## Supplemental methods

### Population Pharmacokinetic Model Simulation

The pediatric population PK model of infliximab by Xiong et al. (4), incorporating time-varying covariates as described by Samuels et al. (14), was implemented using the mrgsolve R package to simulate PK profiles. The two-compartment model included inter-individual variability (IIV) on CL (25.1%) and four key covariates influencing CL: weight (WT), albumin (ALB), erythrocyte sedimentation rate (ESR), and neutrophil CD64 (nCD64) (4). WT was also incorporated for the volumes of the central and peripheral compartments and inter-compartmental clearance, using allometric scaling. Time-varying changes in WT, ALB, ESR, and nCD64 were modeled using Emax functions to capture dynamic responses during infliximab treatment (14). Residual errors were incorporated into the simulations to reflect typical assay variability (CV% = 10%) in infliximab measurements (21). CL values were constrained within the 2.5th and 97.5th percentiles of the simulated distribution to prevent the generation of extreme CL values.

### State, Action, and Q-function Definitions

Each state at decision point *t* (denoted *S*) was defined as a feature vector:

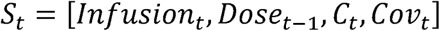

where Infusion_t_ represents the one-hot vector indicating the current infusion number; Dose_t-1_ is a vector representing the prior doses administered at infusion 1 and doses and intervals at infusion 3 (set to zero if the infusion has not yet occurred, e.g., the dose and intervals for infusion 3 are set to zero at the infusion 3 state); C_t_ is the trough concentration prior to the current infusion t (set to zero for infusion 1); and Cov_t_ is a vector of patient-specific covariates (WT, ALB, ESR, nCD64, and ATI) at infusion t. C_t_ and Cov_t_ were assumed to be available at each infusion t. All input variables were log-transformed and scaled to a range of 0 to 1 using min–max normalization.

The action (*A*) was selected from a clinically feasible dosing range, defined as 1–10 mg/kg for infusion 1, and 1–20 mg/kg with dosing intervals of 4–12 weeks for infusions 3–4. These options were represented as discrete integer-valued vectors as follows:

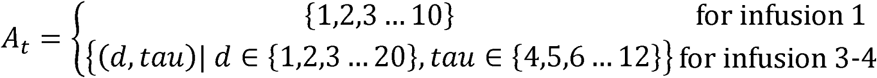

Where d indicates the dose (mg/kg) and tau indicates the dosing interval (weeks).

The Q-function is defined as follows:

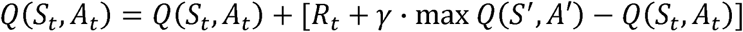

Here, S represents the current state at infusion t, A is the selected action (dose in mg/kg), and R is the reward received at infusion t. The term “max Q(S′, A′)“ denotes the maximum expected future reward at the next state S′, evaluated over all possible actions A′. The discount factor γ represents the relative importance of future rewards and was set to 0.95.

### Implementation of Deep Q-Networks

A Deep Q-Network (17) was implemented in R using the keras3 and tensorflow libraries to predict Q-values for dosing decisions based on the current patient state. The model adopted a dueling network architecture (18) with three shared hidden layers (128 ReLU-activated units each) followed by two separate output heads. The first head predicted Q-values for 10 discrete dose options (1–10 mg/kg) for infusion 1, while the second output Q-values for 180 dosing scenarios representing all combinations of 20 dose levels (1–20 mg/kg) and 9 inter-dose intervals (4–12 weeks) for infusions 3 and 4.

The network was trained using the Huber loss function and the Adam optimizer with an initial learning rate of 0.001 and a decoupled weight decay of 0.01. The learning rate decay followed an exponential schedule, with decay tuned to reach 0.0005 by episode 80,000. A Double DQN (19) strategy was employed to mitigate Q-value overestimation, and a target network was synchronized every 100 episodes.

Training used a prioritized experience replay (PER) buffer (20) with a maximum of 60,000 entries. Experiences were sampled based on rank-based stochastic prioritization, with priority weights derived from temporal difference (TD) error. Importance sampling weights were applied to correct for the bias introduced by prioritization, with PER parameters set to α = 0.6 and an initial β = 0.4, which was linearly annealed to 1.0 over training. The TD error for each experience was updated after every training step.

An ε-greedy exploration policy guided action selection, with ε decaying exponentially from 1.0 to 0.01 using the formula:

ε = max(0.01, ε × (1 - exp(log(0.01)/75000)))

A batch size of 32 was used for training. Target Q-values were computed using the Double DQN formulation and adjusted for each dosing phase. The model was trained for 80,000 episodes, and convergence was monitored every 100 episodes by evaluating the probabilities of target attainment (PTA) at infusions 3, 4, and 5 in a validation cohort of 1,000 virtual pediatric patients. Model hyperparameters were selected based on iterative evaluation of training stability and convergence. All analyses were conducted in RStudio (version 2024.04.0) using R (version 4.4.0) and Rtools 4.4. PK simulations were performed with the mrgsolve package (version 1.5.2). Neural networks were implemented and trained using the TensorFlow (version 2.16.0) and Keras3 (version 1.2.0) packages in R, interfacing with Python 3.11.8 via the reticulate package (version 1.40.0).

---

**Table S1.** Patient characteristics from real-world data used to evaluate the DQN model. The median and range are shown.

|  | Infusion 3 | Infusion 4 | Infusion 5 |
| --- | --- | --- | --- |
| n | 41 | 38 | 35 |
| Age (years) | 14.3 (8.2 - 19.4) | 14.4 (9.0 - 19.4) | 13.7 (4.3 - 19.4) |
| WT (kg) | 39.8 (18.3 - 91.4) | 46.4 (23.9 - 96.3) | 45.2 (15.1 - 102.2) |
| ALB (g/dL) | 3.2 (1.7 - 4.4) | 3.7 (2.6 - 4.8) | 3.7 (3.2 - 4.7) |
| ESR (mm/h) | 18 (2 - 108) | 7.5 (1 - 57) | 9 (1 - 58) |
| nCD64 ratio | 6.5 (2.9 - 18.6) | 4.7 (2.7 - 19.4) | 5 (2.8 - 22.8) |
| Dose at infusion 1-2 (mg/kg) | 6.1 (4.1 - 12.1) | 6.1 (4.1 - 9.5) | 6.0 (4.1 - 12.1) |
| Dose at infusion 3 (mg/kg) | - | 5.9 (3.8 - 12.1) | 5.7 (3.8 - 12.1) |
| Dosing intervals at infusion 3 (weeks) |  | 7.9 (5.1 - 9.1) | 8.0 (5.1 - 8.9) |
| Dose at infusion 4 (mg/kg) | - | - | 5.6 (3.8 - 11.6) |
| Dosing intervals at infusion 4 (weeks) |  | - | 7.6 (4.0 - 13.0) |
| Pre-infusion 3 trough (µg/mL) | 16 (0.4 - 37) | 17.0 (0.4 - 37) | - |
| Pre-infusion 4 trough (µg/mL) | - | 6.5 (0.4 - 29) | 5.0 (0.4 - 23) |
| Pre-infusion 5 trough (µg/mL) | - | - | 4.7 (0.6 - 33) |

## Notes

**Funding:** This work was supported by the National Institute of Diabetes and Digestive and Kidney Diseases at the National Institutes of Health [grant number DK132408], Crohn’s and Colitis Foundation, and the Cincinnati Children’s Research Foundation.

**Conflicts of Interest:** J. S. H. is on advisory boards for Janssen, Abbvie, Lilly, and Genentech and is a consultant for Takeda and Pfizer. The other authors report no conflicts of interest.

### Competing Interest Statement

J. S. H. is on advisory boards for Janssen, Abbvie, Lilly, and Genentech and is a consultant for Takeda and Pfizer. The other authors report no conflicts of interest.

